# Fhod3 in zebrafish supports myofibril stability during growth of embryonic skeletal muscle

**DOI:** 10.1101/2025.05.17.654676

**Authors:** Aubrie Russell, Tracy Eng, Jeffrey D. Amack, David Pruyne

**Affiliations:** Department of Cell and Developmental Biology, State University of New York Upstate Medical University, 750 East Adams St, Syracuse, NY 13210

**Keywords:** skeletal muscle, sarcomere, formin, Fhod3, muscular dystrophy, zebrafish

## Abstract

**Background:** Actin filament organization in cardiomyocytes critically depends on the formin Fhod3, but a role for Fhod3 in skeletal muscle development has not yet been described.

**Results:** We demonstrate here that in zebrafish mutated for one of two *fhod3* paralog genes, *fhod3a*, skeletal muscle of the trunk appears normal through 2 days post-fertilization, but afterward exhibits myofibril damage, including gaps between myofibrils and myofibril fragmentation. Despite the apparent progressive nature of myofibril damage, *fhod3a* mutant embryos differ from muscular dystrophy models in that damage is exacerbated by inhibition of muscle activity, and *fhod3a* mutants show no evidence of sarcolemma disruption. Rather, our results suggest myofibril damage coincides with growth of the skeletal muscle fiber contractile apparatus. We find that neither the second *fhod3* paralog, *fhod3b*, nor the related *fhod1* contribute to embryonic skeletal muscle development, but simultaneous mutation of *fhod3a* and *fhod3b* was associated with pericardial edema suggestive of cardiac dysfunction.

**Conclusions:** Taken together, these results indicate *fhod3-*encoded formins are dispensable for initial myofibril assembly in skeletal muscle, but promote myofibril stability during muscle fiber growth. This is the first demonstration that Fhod3 contributes to skeletal muscle development in a vertebrate.

## INTRODUCTION

Striated muscle cells of vertebrates, which includes cardiac and skeletal muscle fibers, host an elaborate actin-based cytoskeleton of highly ordered units called sarcomeres, as well as a much smaller population of nonsarcomeric actin filaments.^1^ Within the sarcomere, thin filaments are specialized actin filaments anchored at proteinaceous structures called Z-discs at the sarcomere ends. Muscle myosin II-based bipolar thick filaments at the sarcomere center provide force for muscle contraction by pulling thin filaments inward. The force of contraction is propagated longitudinally by the end-to-end attachment of sarcomeres in long chains called myofibrils, and then through adhesion complexes at myotendinous junctions at muscle fiber ends. Force is also transmitted laterally to costameres, which are circumferential bands of adhesion complexes associated with the basement membrane.^2^ Nonsarcomeric actin filaments and desmin-based intermediate filaments likely mediate transmission of force laterally by linking myofibrils to costameres, and linking adjacent myofibrils to each other.^1^

One key question in understanding the organization of the muscle actin cytoskeleton is the identity of factors that control the assembly of the different populations of actin filaments in the muscle cell. Formins are actin-organizing proteins that have been implicated in regulating the actin cytoskeleton in many cell types.^3^ Members of one formin subgroup, the Formin HOmology Domain-2 containing (FHOD) proteins, have been shown to be critical for proper sarcomere organization in several vertebrate and invertebrate striated muscles.^4^ For example, cardiac muscle fibers in *fhod3* knockout mice are able to assemble immature sarcomeres through embryonic day (E) 8.5, but sarcomeric organization is essentially lost by E9.5 and the animals die by E11.5 due to cardiac dysfunction.^5^ Similarly, knockdown of *fhod3* expression through using siRNA or shRNA strongly inhibits formation of mature myofibrils in cultured rat or human induced pluripotent stem cell (hiPSC)-derived cardiomyocytes.^6, 7, 8^ Pointing to the conservation of this function, flies wholly or partially disrupted for expression of their only FHOD homolog, Fhos, are unable to assemble myofibrils in their indirect flight muscle and show cardiac muscle dysfunction, respectively.^9, 10^ In *C. elegans*, absence of its sole FHOD homolog, FHOD-1, also perturbs the muscle cytoskeleton but with the relatively modest effect of slowed sarcomere assembly, and the formation of Z-disc analogs prone to deformation and loss of thin filament attachment during muscle contraction.^11^

The ability of FHOD formins to stimulate actin filament assembly appears critical for their ability to promote sarcomere organization. Functioning as homodimers, formins nucleate new actin filaments through dimerized formin homology-2 (FH2) domains.^12,13, 14^ After filament nucleation, the FH2 dimer remains associated with the fast growing “barbed end” of an actin filament while accommodating the addition of new actin monomers to the growing end, an activity called processive capping.^12, 15, 16^ The FH2 domain/actin interaction is mediated in part through an invariant FH2 domain isoleucine.^14, 17, 18^ Mutation of this residue to alanine diminishes or abolishes formin-stimulated actin filament assembly for many formins, including fly Fhos and mouse Fhod3.^19^ Mutation of this isoleucine in rodent Fhod3, fly Fhos, or worm FHOD-1 all diminish or abolish their ability to promote sarcomere organization in their respective muscles.^6, 10, 11, 20^ Formin-mediated actin filament assembly is also supported by the formin homology-1 (FH1) domain, which recruits the actin monomer-binding protein, profilin, through multiple polyproline stretches.^13, 15, 21^ Rapid transfer of actin monomers from profilin to the FH2-bound barbed end accelerates filament elongation.^22^ Disrupting this mechanism by replacement of the polyproline stretches of human Fhod3 in cultured cells, or by deletion of profilin or the FHOD-1 polyproline stretches from the worm also impairs sarcomere organization.^11, 23^

In the face of conserved use of FHOD formins in striated muscle across many species, and the strong homology between vertebrate cardiac and skeletal muscle sarcomeres, it is striking no role has been described for a FHOD homolog in vertebrate skeletal muscle. Zebrafish provides a useful alternative model to test FHOD formin contributions to skeletal muscle development. Zebrafish skeletal muscle is highly homologous to that of mammals, but more readily observable during embryogenesis by virtue of the fact that fish embryos are fertilized and develop externally, and are optically transparent. Furthermore, the tissues of zebrafish embryos can be directly oxygenated by the surrounding water, allowing embryogenesis to occur up to 5 days post-fertilization (dpf) even in absence of a cardiovascular system.^24^ As skeletal muscle fibers with mature myofibrils are already present in some somites by 24 hours post-fertilization (hpf),^25^ there is an ample window of time to examine skeletal muscle development even if Fhod3 loss results in cardiac dysfunction. Zebrafish encode two *fhod3* paralog genes, *fhod3a* and *fhod3b*, as well as one *fhod1* homolog (Fig.1A). Here we describe the isolation and characterization of fish mutated for *fhod3a* or *fhod3b*, and show that loss of *fhod3a* results in myofibril instability in developing skeletal muscle. This provides the first demonstration that Fhod3 contributes to skeletal muscle development in a vertebrate.

**Figure 1.**
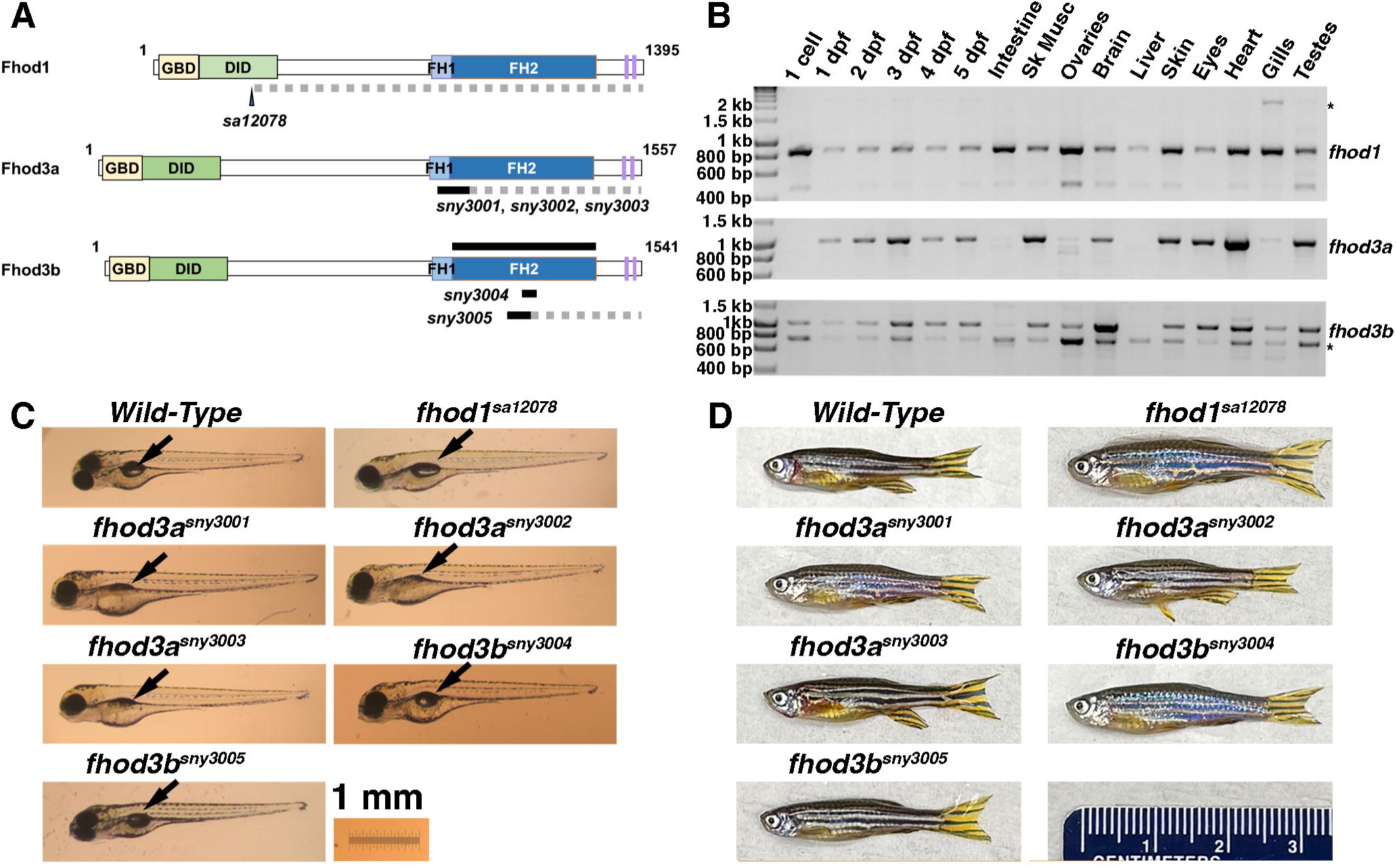
Three *fhod* genes are broadly expressed in zebrafish and individually are not essential. **(A)** The predicted domain organization of three Fhod proteins of zebrafish, showing GTPase-binding domain (GBD), diaphanous inhibitory domain (DID), formin homology-1 (FH1) domain, formin homology-2 (FH2) domain, and two diaphanous autoregulatory domain-like sequences (*purple*). Numbers indicate amino acid residues. Predicted effects of mutant alleles used in this study are shown below each protein. *Black* indicates location of point mutation *sa12078* or sequences deleted by indicated alleles. *Grey hatched lines* indicate regions predicted to be not translated due to premature stop codons and/or shifts in reading frame. (**B)** RT-PCR detection of transcripts for *fhod* genes during embryogenesis and among adult tissues. The identities of DNA bands with gene names were verified by sequence analysis, and bands indicated with asterisks were determined to be non-*fhod* sequences. Results shown are typical for three biological replicates for all samples, except ovaries with two biological replicates, and testes with one biological sample. **(C)** and **(D)** Homozygous maternal/zygotic *fhod* mutants appear morphologically normal as **(C)** 5 dpf embryos and as **(D)** adult females ages 3-10 months, with the exception of a delay in swim bladder inflation (**C**, *arrows*) in *fhod3a* mutant embryos.

## RESULTS

### Three *fhod* genes are broadly expressed in zebrafish

To probe the timing of zebrafish *fhod* gene expression during embryogenesis, we isolated mRNA from whole wild-type embryos and performed reverse transcription-PCR (RT-PCR) (Fig.1, shown n = 1 of 3 biological replicates with same results). Products of the predicted molecular weights were verified through sequence analysis to be *fhod* transcripts (Fig.1B), while additional bands were identified as non-*fhod* sequences (*asterisks*). Transcripts for *fhod1* and *fhod3b* but not *fhod3a* were detected in unfertilized eggs, demonstrating those mRNAs are contributed maternally. All three transcripts were detected in fertilized embryos at 1 to 5 dpf, indicating all three *fhod* genes are transcribed zygotically. A previous high throughput *in situ* hybridization (ISH) screen suggested *fhod1* is broadly expressed throughout embryogenesis,^26^ while two single cell RNA sequencing studies indicate *fhod3a* is primarily expressed in skeletal muscle, cardiac muscle, and the retina during embryogenesis, while *fhod3b* is broadly expressed at low levels.^27, 28^ Unfortunately, our own attempts to probe expression of *fhod3a* and *fhod3b* by ISH yielded weak and inconsistent signals, with similar stain from sense-coding control probes.

To examine the tissue distribution of *fhod* gene expression in adults, we isolated mRNA from select organs from 9-month-old wild-type animals and performed RT-PCR (Fig.1B, shown is 1 of 3 biological replicates with same results, except ovaries as 1 of 2 replicates, and testis as the sole sample). Supporting our observation that *fhod1* and *fhod3b* but not *fhod3a* are maternally provided, we detected transcripts for *fhod1* and *fhod3b* in ovaries, but minimal signal for *fhod3a*. We also observed strong amplification of only *fhod1* and *fhod3b* in the gills, and only *fhod1* in the liver and intestine. For other sampled tissues, skeletal muscle, brain, skin, heart, eye, and testis, we detected transcripts for all three genes. Thus, all three *fhod* genes are expressed during embryogenesis and are broadly expressed among adult tissues.

### The *fhod* genes are not individually essential

To generate fish lacking functional *fhod3a* or *fhod3b*, we performed CRISPR-Cas9-mediated mutagenesis targeting each gene separately, and screened the F2 generation for CRISPR-Cas9-induced deletions. We isolated three alleles for *fhod3a* (*fhod3a^sny3001^*, *fhod3a^sny3002^*, and *fhod3a^sny3003^*) and two alleles for *fhod3b* (*fhod3b^sny3004^* and *fhod3b^sny3005^*). All three *fhod3a* alleles eliminate coding sequence for part of the FH1 domain and the lasso region of the FH2 domain critical for FH2 dimerization, and introduce a premature stop codon (Fig.1A; see Experimental Procedures section for detailed information on alleles). A conceptual translation suggests *fhod3b^sny3004^* produces Fhod3b lacking the L- and M-helices of the FH2 domain, as defined from the Bni1p FH2 domain structure,^14^ while *fhod3b^sny3005^* eliminates FH2 domain helices J through M and introduces a premature stop codon. Due to the protein-coding sequences deleted by each (shown in Fig.1A), we predict any Fhod3 product of the *fhod3a* and *fhod3b* alleles would lack domains critical for formin-mediated actin filament assembly activity. Furthermore, mRNAs from *fhod3a^sny3001^*, *fhod3a^sny3002^*, *fhod3a^sny3003^*, and *fhod3b^sny3005^* are likely to be subject nonsense-mediated mRNA decay (NMD). For analysis of *fhod1* function, Hani Suleiman, MD PhD (Washington University School of Medicine) kindly shared fish bearing *fhod1^sa12078^*, encoding a premature stop codon early in the *fhod1* gene (Fig.1A) isolated by the Stemple Lab (Wellcome Trust Sanger Institute, Cambridge) (Fig.1A). This allele is predicted to truncate most of the Fhod1 protein (Fig.1A), and is also a likely candidate for NMD.

To test whether *fhod1*, *fhod3a*, or *fhod3b* are individually essential for viability, we mated male and female fish that were heterozygous for the same allele, and grew their progeny to adulthood before genotyping. We readily recovered adult homozygotes for each allele. Thus, zygotic expression of no individual *fhod* is essential, although we do not rule out the possibility of a low to moderate level of premature mortality among homozygous mutants. To test whether maternal contribution of any individual *fhod* is essential, we mated male and female fish homozygous for the same allele. Maternal and zygotic null animals for all alleles appeared grossly normal as embryos/young larvae (Fig.1C). The only overt phenotype we observed by simple inspection was that all three *fhod3a* mutant strains consistently exhibited a 1- to 2-day delay in inflating their swim bladder compared to wild-type, *fhod1*, and *fhod3b* embryos (Fig.1C, *arrows*). Moreover, these animals were grossly normal on reaching adulthood (Fig.1D) and were themselves viable, showing there is no maternal requirement for either *fhod3* paralog for viability. Together, our results indicate no single *fhod* gene is essential for viability.

### Loss of *fhod3a* results in myofibril disorganization in developing embryonic skeletal muscle

To examine whether any *fhod* gene contributes to skeletal muscle development, we fixed wild-type and homozygous mutant embryos at 5 dpf, and stained with fluorescently labeled phalloidin to show filamentous actin (F-actin). To avoid the potentially confounding issue of maternal rescue, we examined the progeny of homozygous mutant parents. Skeletal muscle fibers arise in the somites by 24 hpf.^25^ At 5 dpf, wild-type skeletal muscle fibers contain tightly packed F-actin-rich myofibrils that result in a smooth appearance across the muscle fiber (Fig.2A). We observed the same in *fhod1^sa12078^*and *fhod3b^sny3005^* embryos. However, embryos bearing any of the three *fhod3a* mutant alleles displayed unevenness and partial gaps in F-actin along myofibrils (*arrowheads*). In *fhod3a* mutants, many individual myofibrils appeared partially separated from each other (Fig.2B, *arrows*), and obliquely-oriented F-actin strands occupied regions between more widely separated myofibrils (Fig.2B, *arrowheads*). Immunostain showed these strands also contain sarcomeric α-actinin and muscle myosin II (Fig.2B, C, *double arrowheads*), indicating they are fragments of broken myofibrils. Immunostain using F59, an antibody that recognizes muscle myosin II specifically expressed in slow-twitch muscle fibers in zebrafish,^29^ revealed a similar myofibril disorganization in slow-twitch fibers (Fig.2D, *arrowheads*) as seen in non-stained fast-twitch muscle fibers (n = 14 wild-type embryos in five experiments, 510*fhod3a^sny3001^* in two experiments, 7 *fhoda^sny3003^*embryos in three experiments).

**Figure 2.**
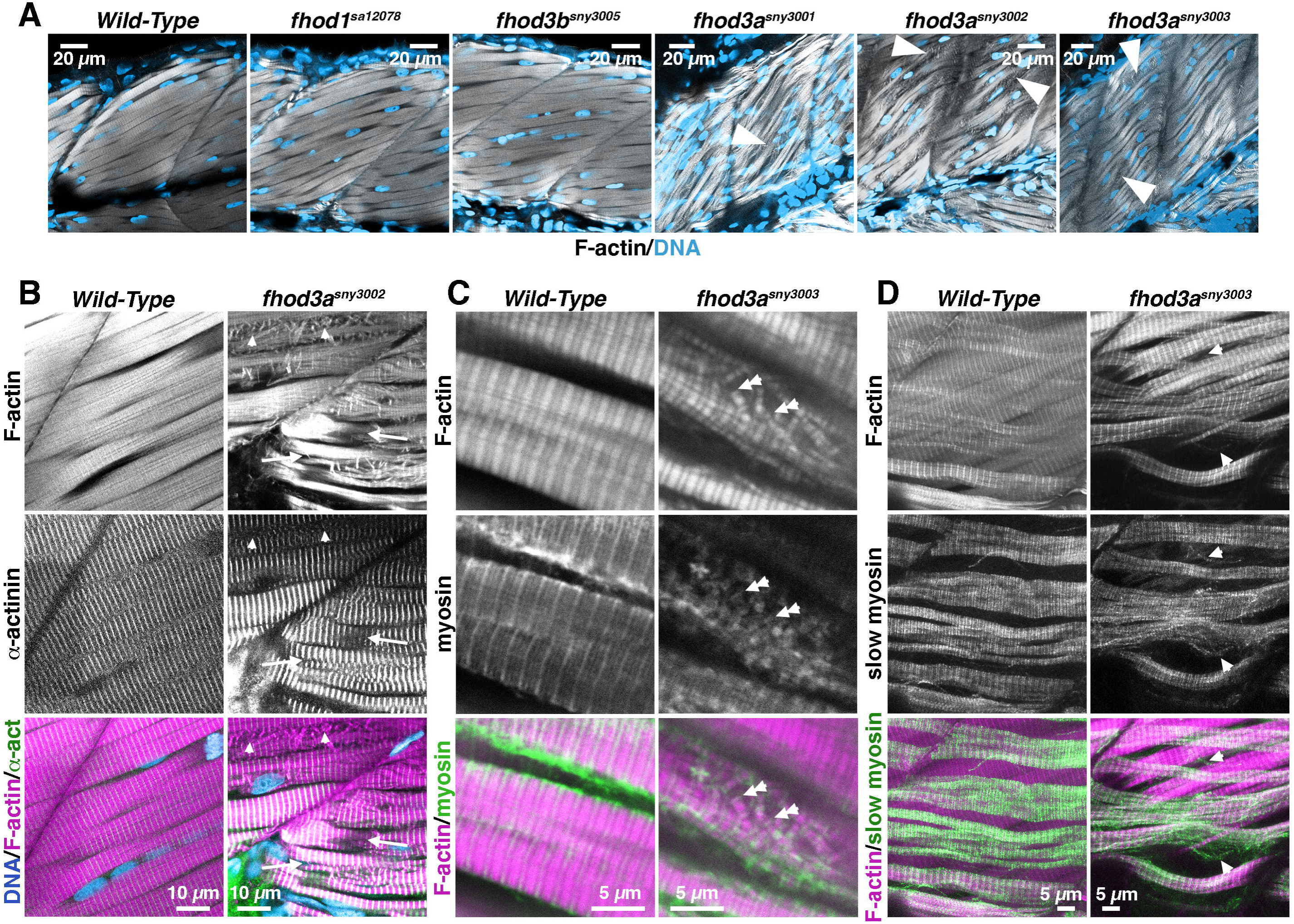
Mutants for *fhod3a* but not *fhod1* or *fhod3b* have broken myofibrils in skeletal muscle during embryogenesis. (**A)** Single plane confocal images of embryos stained at 5 dpf with Alexa-568-phalloidin to show F-actin and DAPI to show nuclei. Arrows indicate F-actin disorganization in *fhod3a* mutants. **(B)** Single plane confocal images of 5 dpf embryos stained for F-actin and sarcomeric α-actinin. Arrows indicate in the *fhod3a* mutant embryo gaps between myofibrils in *fhod3a* mutants. Arrowheads indicate F-actin- and α-actinin-positive myofibril fragments. **(C)** Single plane confocal images of embryos stained at 5 dpf for F-actin and muscle myosin II. Double arrowheads indicate myofibril fragments positive for F-actin and myosin in the *fhod3a* mutant embryo. **(D)** Maximum intensity projections of confocal slices of stained at 5 dpf for F-actin and muscle myosin II specific for slow-twitch muscle (slow myosin). Arrowheads indicate disrupted myofibrils in slow-twitch fibers in the *fhod3a^sny3003^* embryo (results shown are typical for n = 14 wild-type and 7 *fhod3^sny3003^*embryos).

The highly ordered arrangement of myosin filaments in skeletal muscle polarizes light, resulting in birefringence when viewed under polarized light microscopy.^30^ To confirm that the myofibril disruption that we observed was not an artifact of poor fixation, we examined live wild-type and *fhod3a^sny3002^* embryos at 5 dpf using polarized light microscopy (Fig.3A). Wild-type embryos showed an even level of birefringence in somites. observed uneven birefringence in live 5 dpf *fhod3a^sny2003^* embryos under polarized light, as compared to a more even level of birefringence in somites of wild-type embryos (Fig.3A). This was reflected in the intensity of the birefringence signal along line scans in wild-type embryos, which showed relatively constant levels punctuated by crisp reductions marking the septa between somites (Fig.3B). The overall level of birefringence was in *fhod3a^sny3003^* embryos was similar to wild-type, but the distribution was more variable (Fig.3A). This was reflected in a range of patterns seen in line scans. For some *fhod3a^sny3003^* embryos, the pattern of birefringence resembled wild-type (e.g., dorsal line scans of *fhod3a^sny3003^*fish 3 and 4, and ventral line scans fish 5 and 7), while in others, birefringence increased from the anterior toward the posterior border of the somite (e.g., dorsal line scans *fhod3a^sny3003^* fish 2, 5, and 6, and ventral line scans fish 1 and 4). In others, birefringence was sufficiently variable that the positions of somite boundaries were not obvious in line scans (e.g., dorsal line scan *fhod3a^sny3003^* fish 1, and ventral line scans fish 2 and 3), although septa were still identifiable in images. These results suggest that while the overall amount of ordered myofilaments is not grossly reduced in *fhod3a* mutant embryonic muscle, the distribution of myofilaments is altered, as might be expected from myofibril fragmentation.

**Figure 3.**
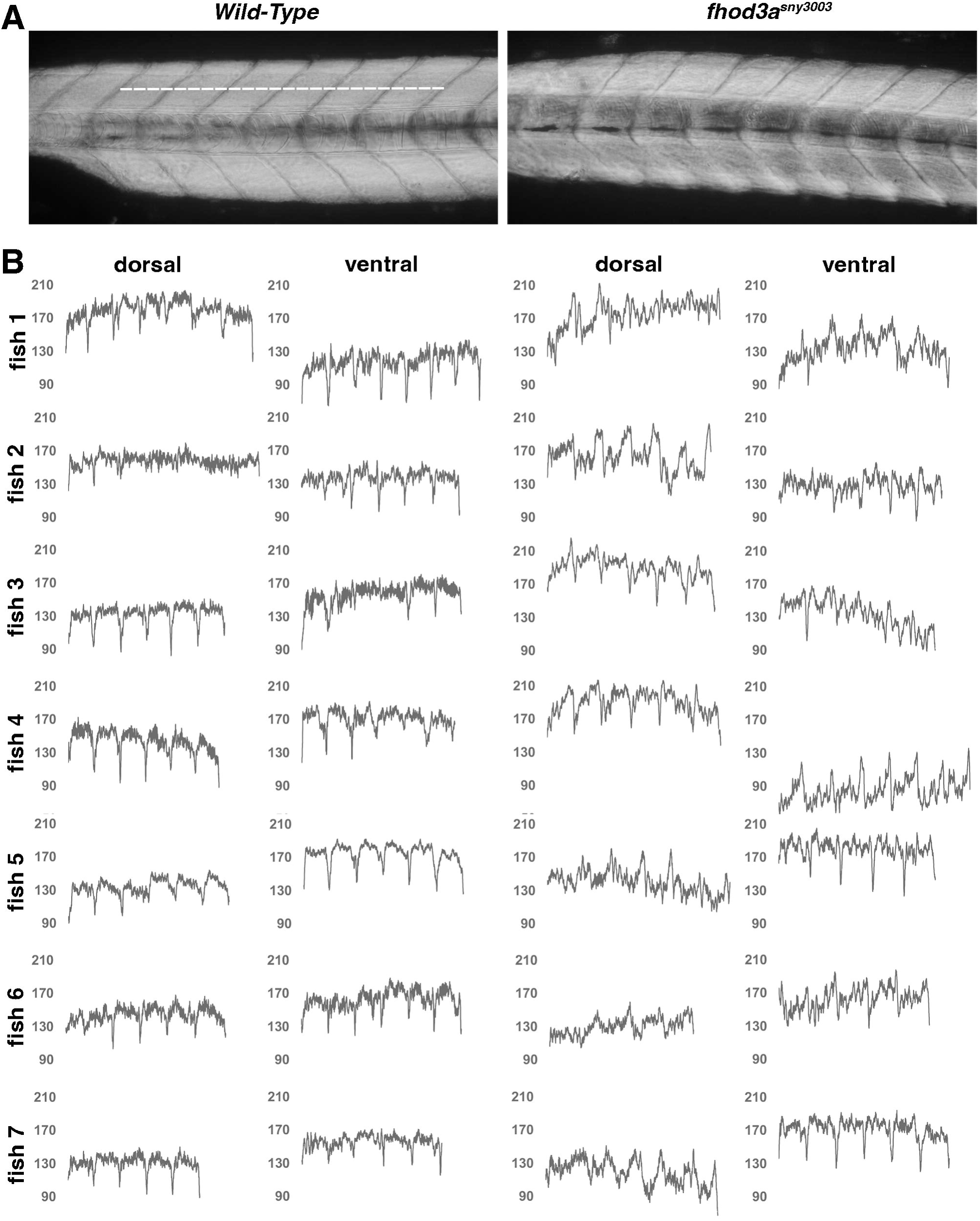
Polarized light microscopy shows myofilament disorganization in live *fhod3a* mutant embryos. **(A)** Live embryos were viewed at 5 dpf using polarized light microscopy. Shown for the wild-type embryo is a representative line over which birefringence was measured to generate line scans as shown in **B**. (**B)** Birefringence was measured in 5 dpf embryos along two lines spanning six somites, one positioned midway between the notochord and the dorsal somite boundary (as shown in **A**), and one positioned between the notochord and the ventral somite boundary. Birefringence intensity values are reported on the vertical axis.

To determine whether *fhod3a* mutants are defective in initial myofibril assembly or in later growth, we examined somites at earlier stages of development. At 2 dpf, skeletal muscle fibers in all three *fhod3a* mutant strains were similar to wild-type, with myofibrils containing striated F-actin and well-organized Z-lines (Fig.4A). At 3 dpf, where wild-type myofibrils were smooth in appearance, those of all three *fhod3a* mutant strains appeared rougher, with F-actin strands partially dissociated from myofibrils (Fig.4A, *arrows*). In single confocal sections, F-actin disorganization appeared more prevalent among muscle fibers in anterior somites, where ∼ 62 - 86% of myofibrils showed F-actin gaps or misplaced strands compared to ∼ 13 - 43 % of myofibrils of the tail (Fig.4B). However, the true proportion of damaged myofibrils was likely higher in both regions, as additional damage was frequently observed in higher or lower focal planes. At 5 dpf myofibrils, the proportion of visibly damaged myofibrils in single confocal sections was similar in each *fhod3a* mutant strain (to ∼ 87-88% of myofibrils near the head and 16 - 32% of myofibrils in the tail; Fig.4B), but the damage was qualitatively greater than at 3 dpf. All three strains had abundant misoriented F-actin strands (Fig.4A, *arrows*) and regions where myofibrils appear to have been disrupted or depleted for F-actin (*arrowheads*). Thus, absence of *fhod3a* function does not prevent initial myofibril assembly in skeletal muscle, but results in apparent myofibril instability during muscle fiber growth.

**Figure 4.**
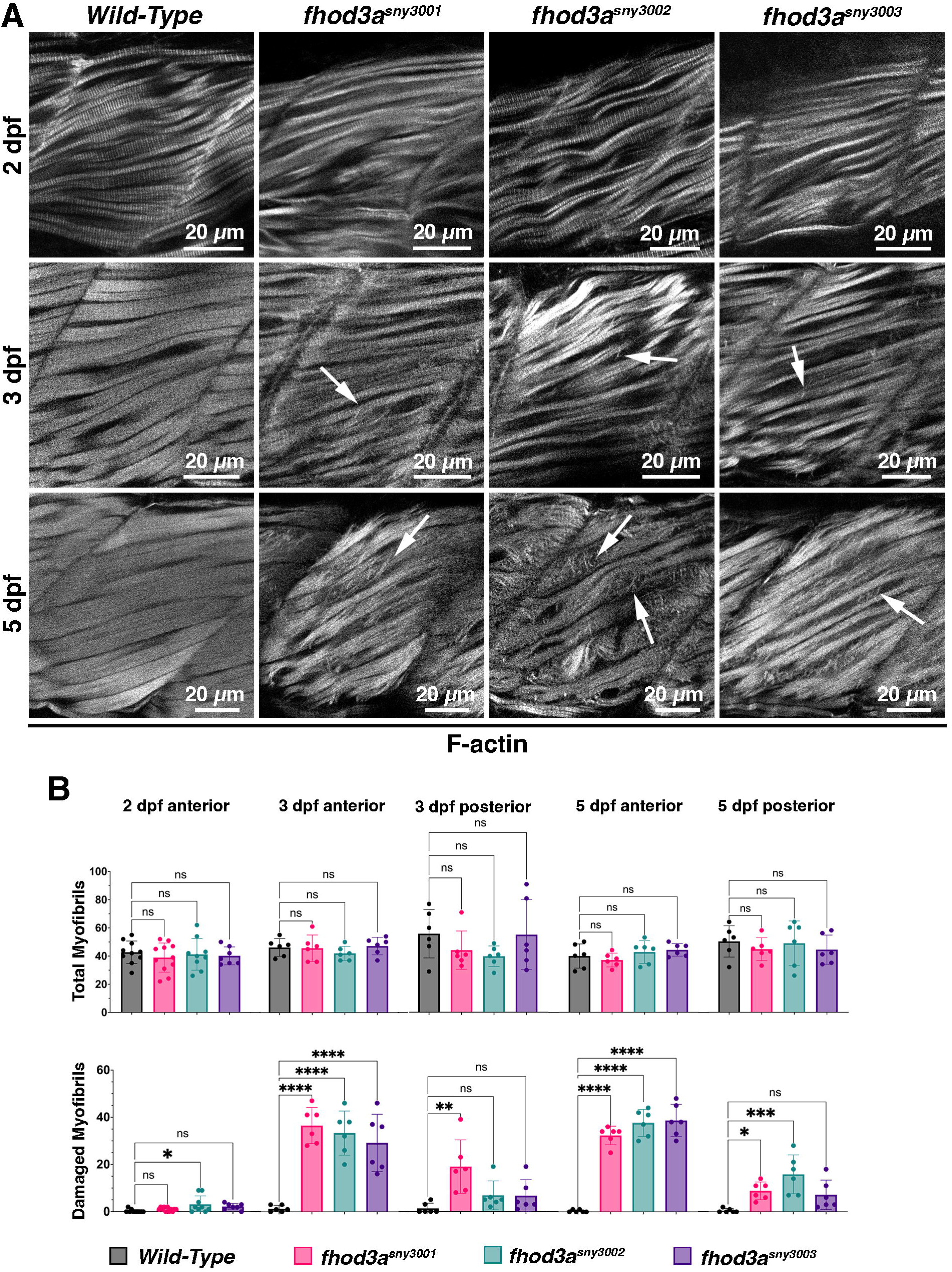
Myofibril damage in *fhod3a* mutants develops later in embryogenesis. **(A)** Single plane confocal images of anterior somites in embryos stained for F-actin. Arrows indicate partially displaced F-actin strands in *fhod3a* mutant embryos older than 2 dpf. Arrowheads indicate regions of myofibril disruption in 5 dpf *fhod3a* mutant embryos. **(B)** The presence or absence of myofibril damage was quantified from confocal images acquired at three different focal planes of an anterior somite (somite 5) in 2, 3, and 5 dpf embryos stained for F-actin, and confocal images from three different focal planes of a posterior somite (penultimate tail somite) in 3 and 5 dpf embryos. Data shown are mean ± SD with individual counts, for n ≥ six fish of each genotype and age. (*) indicates *p* ≤ 0.05; (**) *p* ≤ 0.005; (***) *p* ≤ 0.0005; (****) *p* ≤ 0.0001.

### *fhod3a* skeletal muscle defects do not phenocopy muscular dystrophies

Progressive muscle damage is one hallmark of muscular dystrophies.^31^ We tested whether *fhod3a* mutants exhibit additional phenotypes found in zebrafish models for muscular dystrophy. Muscle activity exacerbates muscle damage in clinical cases of muscular dystrophy,^31,32, 33^ while inhibition of muscle activity reduces muscle damage during embryogenesis in multiple zebrafish muscular dystrophy models.^34, 35, 36^ Expression of the nicotinic acetylcholine receptor inhibitor α-bungarotoxin (α-BTX) is an effective method for long-term immobilization of zebrafish embryos with few side effects.^37^ We injected wild-type, *fhod3a^sny3002^*, and *fhod3a^sny3003^*1-cell embryos with mRNA encoding α-BTX. Injected embryos that remained fully paralyzed at 3 dpf were fixed and stained for F-actin, along with uninjected control embryos. Myofibril organization was unchanged in α-bungarotoxin (α-BTX)-injected wild-type embryos compared to uninjected controls, showing that on its own, α-BTX does not visibly perturb skeletal muscle myofibrillogenesis (Fig.5A) (n = 12 uninjected and 19 injected wild-type embryos, over three experiments). Surprisingly, α-BTX-injected *fhod3a^sny3002^* and *fhod3a^sny3003^* embryos showed a visible increase in the prevalence of misoriented F-actin strands compared to uninjected controls (Fig.5A, *arrows*) (n = 3 uninjected and 3 injected *fhod3^sny3002^*embryos from one experiment, and 13 uninjected and 13 injected *fhod3^sny3003^* embryos, over two experiments). These results suggest that the development of myofibril defects in *fhod3a* mutants is partially countered by muscle activity.

**Figure 5.**
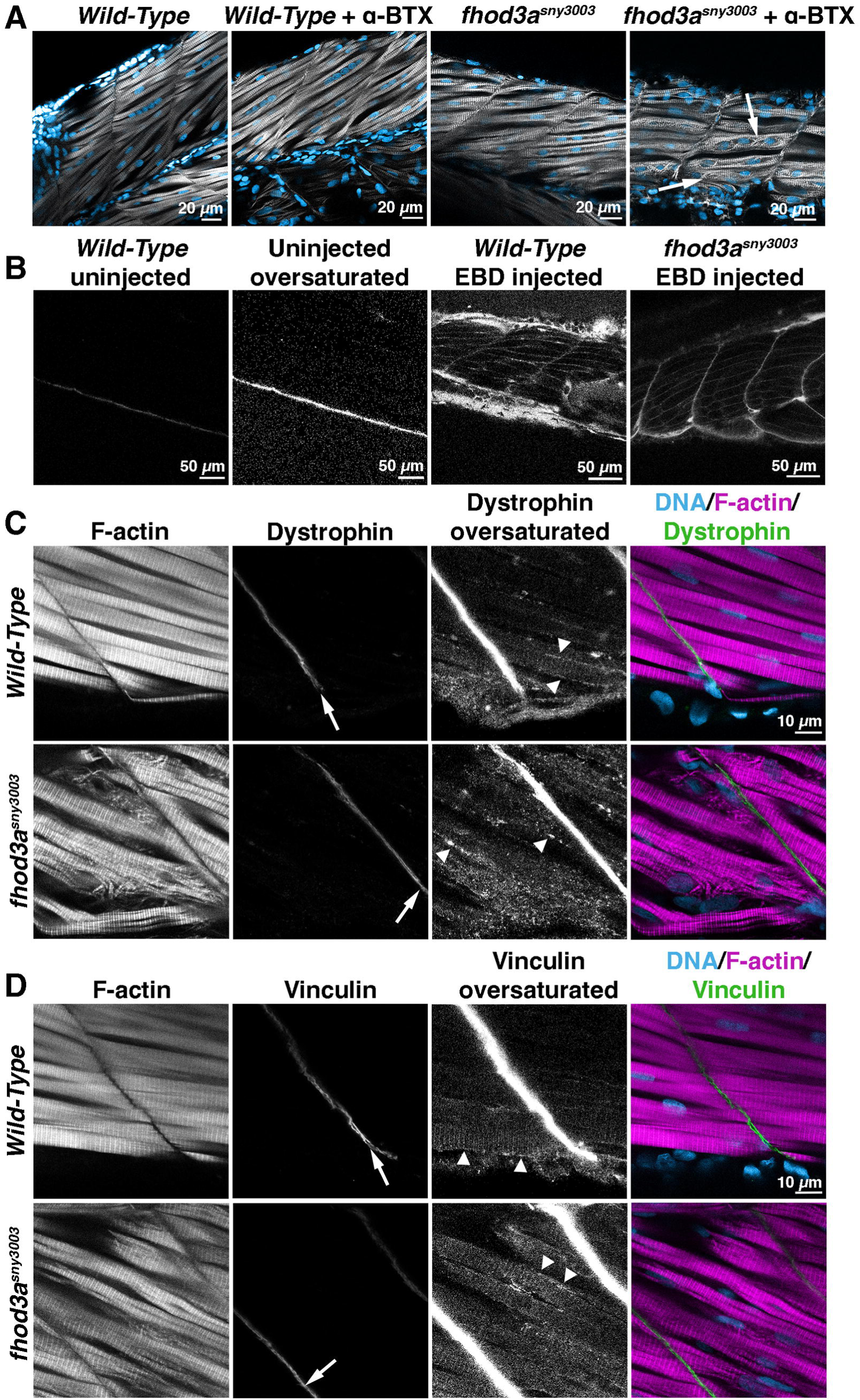
Mutants for *fhod3a* do not share phenotypes found in models for muscular dystrophy. **(A)** Single plane confocal images of embryos stained for F-actin (*white*) and nuclei (*blue)* at 3 dpf. Embryos either were or were not injected with mRNA to express α-BTX to induce skeletal muscle paralysis. Arrows indicate increased F-actin disorganization in the *fhod3a^sny3003^* embryo (results typical for n = 12 to 19 embryos each condition). **(B)** Single plane confocal images of embryos injected or uninjected with Evans blue day (*EBD*) at 5 dpf, and viewed for red fluorescence. Uninjected embryo fluorescence is shown at normal and oversaturation to demonstrate lack of signal compared to signal the vasculature of injected embryos. Note the lack of muscle fiber stain in either strain (results typical for n = 5 embryos each condition). **(C)** and **(D)** Single plane confocal images of embryos stained at 5 dpf to show F-actin, DNA, and either **C** dystrophin or **D** vinculin. Arrows indicate strong presence at myotendinous junctions, and arrows indicate weaker association with costameres (results typical for **C**, n = 9 wild-type, 4 *fhod3a^sny3003^* embryos; **D**, n = 12 wild-type, 6 *fhod3a^sny3003^* embryos).

Loss of sarcolemma integrity is another hallmark of muscular dystrophies, as has been demonstrated in mouse and zebrafish models through the use of Evans blue dye.^38,39,40^ After injection into the vasculature of the zebrafish embryo, Evans blue is unable to cross the sarcolemma and is excluded from skeletal muscle fibers in wild-type animals, whereas in *dmd^sapje^* embryos, the zebrafish model for Duchenne muscular dystrophy, the dye strongly accumulates in muscle fibers by 4-6 hr after injection due to breaks in the sarcolemma.^40^ To test sarcolemmal integrity in a *fhod3a* mutant, we injected Evans blue dye into the vasculature of wild-type and *fhod3^sny3003^* embryos at 5 dpf, when myofibril damage is pronounced in the mutant. After 4.75 hr of recovery, the dye was visible in the vasculature between muscle fibers and somites in wild-type and *fhod3a^sny3003^*, but was not visible within skeletal muscle fibers for either strain (n = 5 injected and uninjected embryos of each genotype) (Fig.5B). Thus, *fhod3a* mutants show no indication of sarcolemma disruption.

Finally, many muscular dystrophies result from disturbances of cell adhesion complexes associated with costameres and myotendinous junctions, particularly the dystrophin-glycoprotein complex (DGC).^31,41^ To examine whether *fhod3a* mutations result in gross disturbance if the DGC, we immunostained embryos for the DGC component, dystrophin. At 5 dpf, dystrophin is strongly associated with myotendinous junctions in wild-type embryos (Fig.5C, *arrows*), while association with costameres is patchy (*arrowheads*) but increases later in development.^35^ We observed the same in *fhod3a^sny3003^*embryos at 5 dpf (Fig.5C), suggesting the DGC is not grossly perturbed for localization (n = 9 wild-type embryos in three experiments, 4 *fhod3a^sny3003^*embryos in two experiments). Integrin-based complexes containing vinculin and talin also associate with costameres and myotendinous junctions.^1^ Similar to dystrophin, we observe strong association of vinculin with the myotendinous junctions (Fig.5D, *arrows*) in wild-type, *fhod3a^sny3002^*, and *fhod3a^sny3003^*embryos at 4 and 5 dpf, and weaker association with costameres (*arrowheads*) (n = 12 wild-type embryos in four experiments, 6 *fhod3a^sny3002^* embryos in two experiments, 6 *fhod3a^sny3003^* embryos in one experiment). In summary, by several criteria, *fhod3a* mutants do not resemble canonical muscular dystrophy models.

### *fhod3b* plays no role in skeletal muscle, but *fhod3a* and *fhod3b* may play overlapping roles in cardiac muscle

To test whether the relatively mild skeletal muscle phenotype of *fhod3a* mutants is due to partial compensation by *fhod3b*, we crossed *fhod3a* and *fhod3b* mutants and examined their progeny for enhanced skeletal muscle defects. Mating of *fhod3a^syn3002^*homozygotes with *fhod3b^syn3004^* homozygotes yielded normal-appearing doubly heterozygous *fhod3a^syn3002/+^ fhod3b^syn3004/+^*F1 progeny. These F1 fish were then mated to each other, and their F2 progeny were fixed at 5 dpf and stained for F-actin. After microscopic analysis, embryos were genotyped (Table 1). Among the F2 animals, 61 of 73 appeared normal, and 12 had anterior somitic myofibril disorganization similar to that of *fhod3a* homozygous mutants. Strikingly, all 12 embryos with damaged myofibrils were homozygous for *fhod3a^sny3002^*while only one embryo with normal-appearing myofibrils was homozygous for that allele. Thus, there was a significant correlation between *fhod3a^sny3002^*homozygosity and myofibril disorganization (χ^2^ = 66.28, df = 1, *p =* 3.9 × 10^-16^). In contrast, there was no significant correlation between *fhod3b^sny3004^*homozygosity and that phenotype (χ^2^ = 0.77, df = 1, *p* = 0.36). Additionally, these results showed that any putative maternal contribution of wild-type *fhod3a* function from the heterozygous F1 parent does not contribute to myofibril stability.

**Table 1.** Only *fhod3a* and not *fhod3b* affects myofibril stability in skeletal muscle.

| Genotype<sup>a</sup> | Normal myofibrils | Disorganized myofibrils |
| --- | --- | --- |
| *fhod3a*<sup>+/+</sup> *fhod3b*<sup>+/+</sup> | 17 | 0 |
| *fhod3a*<sup>+/+</sup> *fhod3b*<sup>+/-</sup> | 15 | 0 |
| *fhod3a*<sup>+/+</sup> *fhod3b*<sup>-/-</sup> | 6 | 0 |
| *fhod3a*<sup>+/-</sup> *fhod3b*<sup>+/+</sup> | 11 | 0 |
| *fhod3a*<sup>+/-</sup> *fhod3b*<sup>+/-</sup> | 8 | 0 |
| *fhod3a*<sup>+/-</sup> *fhod3b*<sup>-/-</sup> | 3 | 0 |
| *fhod3a*<sup>-/-</sup> *fhod3b*<sup>+/+</sup> | 1 | 6 |
| *fhod3a*<sup>-/-</sup> *fhod3b*<sup>+/-</sup> | 0 | 3 |
| *fhod3a*<sup>-/-</sup> *fhod3b*<sup>-/-</sup> | 0 | 3 |

<sup>a</sup> *fhod3a*<sup>-</sup> indicates *fhod3a*<sup>sny3002</sup>. *fhod3b*<sup>-</sup> indicates *fhod3b*<sup>sny3004</sup>.

In addition to skeletal muscle defects, we anecdotally observed cardiac edema in some F2 embryos at 3 dpf. Although cardiac edema is not a definitive indicator of heart dysfunction, it is frequently associated with defects in genes related to heart or cardiovascular function.^42^ To test whether this phenotype correlated with double homozygosity for *fhod3a* and *fhod3b*, we mated male and female fish of the genotype *fhod3a^sny3002/+^ fhod3b^sny3004/sny3004^*, and examined their progeny. The majority of embryos from one cross had a noticeable developmental delay at 24 hpf, with 28 of 49 embryos having 3 to 10 somites (∼ 12 hr delay), and the remaining 21 having 18 to 21 somites (∼ 5 hr delay). By 48 hpf, all embryos looked mildly underdeveloped, but otherwise grossly normal. By 3 dpf, mild cardiac edema developed in some embryos, followed by significant cardiac edema at 5 dpf. Genotyping showed that 13 of the 16 edematous embryos from this cross were *fhod3a^sny3002^ fhod3b^sny3004^* double homozygotes, while only 3 of 33 non-edematous embryos were double homozygotes (Table 2). Thus, there was a strong correlation between cardiac edema and *fhod3a^sny3002^* homozygosity in the *fhod3b^sny3004^* background (χ^2^ = 20.4, df = 1, *p* = 6.35 x 10^-6^). In contrast, we do not observe cardiac edema among the progeny of mated heterozygous *fhod3a^sny3002/+^* fish in a wild-type *fhod3b* background. In summary, *fhod3b* appears to play no role in myofibril stability of embryonic skeletal muscle, but *fhod3a* and *fhod3b* may play overlapping roles in development, and particularly in cardiac development during embryogenesis.

**Table 2.** Mutation of *fhod3a* and *fhod3b* results in cardiac edema at 5 dpf.

| Genotype<sup>a</sup> | No cardiac edema | Cardiac edema |
| --- | --- | --- |
| *fhod3a*<sup>+/+</sup> *fhod3b*<sup>-/-</sup> | 11 | 2 |
| *fhod3a*<sup>+/-</sup> *fhod3b*<sup>-/-</sup> | 19 | 1 |
| *fhod3a*<sup>-/-</sup> *fhod3b*<sup>-/-</sup> | 3 | 13 |

<sup>a</sup> *fhod3a*<sup>-</sup> indicates *fhod3a*<sup>sny3002</sup>. *fhod3b*<sup>-</sup> indicates *fhod3b*<sup>sny3004</sup>.

## DISCUSSION

The formin Fhod3 has been shown to be critical to myofibril organization in cardiac muscle fibers in humans, mice and rats,^5,6,7,8^ but evidence for a role in skeletal muscle has been absent. Using the zebrafish model, we provide the first demonstration that Fhod3 also contributes to skeletal muscle development in a vertebrate.

### Redundancy and uniqueness of zebrafish *fhod3* paralogs

Most vertebrates, including mammals, have two genes that encode FHOD formin family proteins, *fhod1* and *fhod3*, while teleosts such as zebrafish have three, *fhod1* and two *fhod3* paralogs, *fhod3a* and *fhod3b*. In the mouse, *fhod1* is widely expressed but not required for normal development. We find here that *fhod1* is also not essential in zebrafish, based on a premature stop codon-containing allele. In contrast, mouse *fhod3* has more a more restricted expression during embryogenesis, and is essential in multiple tissues during embryogenesis and perinatal development, including the neural plate and heart.^5,43,44, 45^ Surprisingly, we find that neither *fhod3* paralog of zebrafish is essential, based on FH2 domain-disrupting deletion alleles that we have isolated. We suggest that a likely explanation for this difference between fish and mouse is redundancy between *fhod3a* and *fhod3b* for important functions. In a survey of adult zebrafish tissues, we observe that *fhod3a* and *fhod3b* are both expressed in many tissues, including in heart and brain (Fig.1B). While we were not able to determine expression patterns for *fhod3a* or *fhod3b* in the embryo through ISH, data reported from two single cell RNA sequencing (scRNA seq) studies of zebrafish embryos show both genes are expressed in embryonic cardiomyocytes.^27, 28^ In support of their functional redundancy in cardiac muscle, previous simultaneous targeting of *fhod3a* and *fhod3b* for CRISPR-Cas9-mediated mutagenesis resulted in increased incidence of cardiac dysfunction.^46^ Similarly, we observed here that fish doubly homozygous for loss-of-function alleles for *fhod3a* and *fhod3b* exhibit early signs of cardiac dysfunction during embryogenesis, but observed the same phenotype at much lower frequency in embryos with one or more wild-type copies of *fhod3a* or *fhod3b* (Table 2). We anticipate double *fhod3a fhod3b* mutants might serve as a useful model for future study of *fhod3* dysfunction in cardiac muscle.

The two zebrafish embryo scRNA seq studies also reported differences in expression of *fhod3a* and *fhod3b* in other embryonic tissues.^27, 28^ In particular, both studies showed embryonic expression of *fhod3a* is strongest in skeletal muscle, including fast- and slow-twitch fibers, followed by lower expression in cardiac muscle and eye photoreceptors. In contrast, *fhod3b* is expressed at low levels across most tissues of the embryo, with strongest expression in cells of the retina, including photoreceptors. The strong expression of *fhod3a* in skeletal muscle compared to *fhod3b* may explain why we observe that *fhod3a* and not *fhod3b* contributes to myofibril stability during embryogenesis (Fig.2A, Table 1), while our observation that *fhod3b* is expressed with *fhod3a* in skeletal muscle of adult fish (Fig.1B) might explain why adult homozygous *fhod3a* mutants have no overt movement defects.

### *fhod3a* supports myofibril integrity during skeletal muscle fiber growth

Zebrafish skeletal muscle progresses through similar stages of myofibrillogenesis as seen in mammalian and avian cardiac and skeletal muscle: immature sarcomeres are initially organized into premyofibrils, which grow and coalesce into nascent and then mature myofibrils.^47^ This occurs very early in development, such that by 24 hpf skeletal muscle fibers in anteriorly positioned somites (which form earliest in development) already contain mature myofibrils.^47^ This initial myofibril formation appears normal in *fhod3a* mutants, as at 2 dpf they are virtually indistinguishable from wild-type embryos (Fig.4A). Continuing beyond 2 days, muscle fibers in wild-type embryos undergo hypertrophy, including a dramatic accumulation and dense packing of actin filaments into growing myofibrils (Fig.4A, *3 dpf*, *5 dpf*). In contrast, F-actin accumulation is reduced in skeletal muscle fibers of *fhod3a* mutant embryos (Fig.4A). Myofibrils appear thinner (Fig.4A, *5 dpf*) with displaced F-actin strands (Fig.4A, *arrows*). Close inspection shows these strands are sarcomeric in nature, and likely reflect fragmentation of myofibrils (Fig.3B,C, *arrowheads*, *double arrowheads*). Moreover, gaps are often apparent between myofibrils of *fhod3a* mutant skeletal muscle at 5 dpf (Fig.3B, *arrows*).

The relatively late onset of the *fhod3a* mutant phenotype seemed suggestive of a progressive myopathy, similar to what is observed for muscular dystrophies. However, *fhod3a* mutants lack other hallmarks found with other zebrafish muscular dystrophy models. In many clinical and animal models of muscular dystrophy, defective cell adhesions found at costameres and myotendinous junctions underlie dysfunction.^40, 41, 48^ But we observed no evidence of detachment of muscle fibers from myotendinous junctions. Nor did we observe evidence of sarcolemma disruption indicative of costamere dysfunction (Fig.5B), or mislocalization of costamere/myotendinous junction proteins dystrophin or vinculin (Fig.5C, D). Also unlike muscular dystrophies, muscle damage in *fhod3a* mutants was not activity-dependent (Fig.5A). Notably, inhibition of muscle activity prevents damage in zebrafish models of muscular dystrophy, but myofibril damage,^34, 35, 36^ but myofibril damage was exacerbated at 3 dpf in *fhod3a* mutant embryos that were paralyzed (compare Fig.5A, *fhod3a* + α-BTX to Fig.4A, 5 dpf *fhod3a* mutants).

Based on the timing of onset of the phenotype, we suggest Fhod3a activity is important for maintaining myofibril stability during muscle fiber growth. Consistent with this, muscle fibers of the tail, which grow less than those in anterior somites, show more moderate myofibril damage. Interestingly, in the *fhod3* knockout mouse, embryonic cardiomyocytes assemble sarcomeres into premyofibrils that produce a heartbeat, indicating initial sarcomere formation in the mammalian heart also does not require Fhod3.^5^ Loss of sarcomere organization in the *fhod3* knockout heart only occurs over the following days, a period when wild-type cardiomyocytes undergo hypertrophy, with coalescence of premyofibrils into nascent and mature myofibrils.^5^ Thus, a similar mechanism may underlie the role of these formins in both muscle types, although cardiac muscle has a stronger reliance on Fhod3.

The timing of Fhod3a function in zebrafish skeletal muscle development may also explain why a role for Fhod3 in skeletal muscle has not been previously described in mouse. Fhod3 protein is present in mouse embryonic skeletal muscle,^49^ and the *fhod3* gene co-expresses with genes involved in early myofibrillogenesis in a subset of skeletal muscle fiber nuclei in P21 mice, ^51^ but two studies failed to observe skeletal muscle defects related to *fhod3* mutations. An induced knockout of *fhod3* in mouse cardiac and skeletal muscle during perinatal development show cardiac muscle defects at postnatal day 6, but no overt sarcomere defects in the quadriceps (a skeletal muscle).^50^ However, a key caveat to this finding is the timing and extent of Fhod3 protein depletion from the quadriceps could not be determined due to low abundance of Fhod3. Alternatively, mice with a global *fhod3* knockout but temporarily rescued for viability through transgenic expression of *fhod3* in cardiac muscle are expected to be fully lacking Fhod3 in developing skeletal muscle. In such mice, skeletal muscle of the tongue appears normal at embryonic day 19.5, showing Fhod3 is not required for initial myofibrillogenesis in that skeletal muscle.^49^ However, our results suggest Fhod3 might be important for longer term maintenance of myofibrils, particularly during muscle growth.

The *fhod3a* mutations examined here are likely to be null alleles due to NMD. Therefore, we do not definitely attribute Fhod3a function to any particular molecular interaction or activity. However, other models for Fhod3 function in muscle depend on the ability of the formin to promote actin filament assembly.^6, 10, 11, 20^ The actin filament product for Fhod3 in these systems is often presumed to be thin filaments. The strongest evidence for this comes from *Drosophila*, where the Fhod3 homolog, Fhos, assembles new thin filaments and elongates the barbed ends of existing thin filaments as myofibrils thicken.^10^ A similar role for Fhod3a in skeletal muscle might explain the relative paucity of F-actin in myofibrils of embryonic skeletal muscle in *fhod3a* mutants. An alternative speculative model is that Fhod3a might polymerize nonsarcomeric actin filaments composed of cytoplasmic γ-actin that are present between myofibrils and between peripheral myofibrils and costameres.^52, 53^ In mouse skeletal muscle, such nonsarcomeric actin filaments are known to organize the sarcoplasmic reticulum between myofibrils,^53^ and mice with a skeletal muscle-specific knockout of γ_cyto_-actin develop a progressive myopathy likely related to costamere dysfunction.^54, 55^ Additionally, knockdown studies in primary mouse skeletal myotubes has shown γ_cyto_-actin is also critical to maintaining desmin-based linkages between adjacent myofibrils.^56^ This inter-myofibril cohesion, together with myofibril contraction, are critical to pushing nuclei to the muscle fiber periphery.^56^ Such a role might explain why myofibril defects in *fhod3a* mutants are exacerbated by inhibition of muscle activity by α-BTX, particularly in the vicinity of muscle fiber nuclei (Fig.5A). The question of whether *fhod3a* mutants shows defects specifically in organization of nonsarcomeric actin filaments or sarcomeric thin filaments in skeletal muscle will be a key subject of future studies.

## EXPERIMENTAL PROCEDURES

### Zebrafish

All animal care and experiments were approved by the SUNY Upstate Medical University IACUC under protocol #363. Embryos were raised at 28°C. The TAB zebrafish strain was used as wild-type for this study. Fish bearing *fhod1^sa12078^* were isolated by the Stemple Lab (Wellcome Trust Sanger Institute, Cambridge) and kindly shared with us by Hani Suleiman, MD/PhD (Washington University School of Medicine, St. Louis, MO). *fhod1^sa12078^* was verified by Sanger sequence analysis after PCR of *fhod1* using the following primers (Integrated DNA Technologies, Coralville, IA):

fhod1sa12078TestRor: GGC GTA CAT ATC CAA GAA AGG TT

fhod1sa12078TestRev: GGA CGA TCC TTA TAG ACA TTC TGT CT

### RT-PCR

To harvest RNA from adult tissues, four wild-type animals (two males, two females) were euthanized by submersion in ice-cold water, and individual organs were dissected directly into PureZOL (Bio-Rad, Hercules, CA, Cat #7326890) and homogenized with a 28-gauge syringe.

Three biological replicates of each organ were analyzed (from two females and one male), except ovaries (two biological replicates from two female fish), and testes (one testis from one male fish). To harvest embryonic RNA, embryos of each age category were collected from three separate spawns treated as biological replicates. 24 hpf, 2 dpf, and 3 dpf embryos were dechorionated before RNA isolation. Whole embryos were homogenized in PureZOL with a 28-gauge syringe, subject to chloroform extraction, digested with DNAse I (New England Biolabs, Ipswitch, MA, Cat #M0303S), and subject to a second chloroform extraction. 1 µg of RNA was reverse-transcribed using a Superscript 2 First-Strand Synthesis Kit (Invitrogen, Waltham, MA, Cat #11904018) following the manufacturer’s protocol. 1 µL of cDNA was amplified using Taq DNA polymerase (New England Biolabs, Cat #M0273L), and the following primers (Integrated DNA Technologies):

zfhod1cDNAfor_158: TCG TTG TGA CCA CGG ATA AGC

zfhod1cDNArev_1013: ATC AGG AAG TGC TGC CAG AG

zfhod3a_for: GGT TCC GGA GGT GAA GGA CAC

zfhod3a _rev: CAG TCG GGA CGT CCT GTT ACA C

zfhod3b _for: GTC AGT GAC AGC AAC TCA GGT C

zfhod3b _rev: CCA GAA TGA TGA TCT CCT GAC GC

### CRISPR-Cas9 mutagenesis

Strategies for CRISPR-Cas9-mediated mutagenesis of *fhod3a* and *fhod3b* were designed in collaboration with InVivo Biosystems (Eugene, OR). Reagents for mutagenesis were provided by InVivo Biosystems. One-cell embryos were injected with one of two custom injection mixes (InVivo Biosystems). One mix contained 0.5 mg/ml Cas9, two sgRNAs targeting *fhod3a* exons 19 and 20 (of predicted splice isoform X1, Accession XM_068214113, version 1), and oligodeoxynucleotide (ODN) for homology-directed repair (HDR) resulting in 1635 bp in-frame deletion in *fhod3a*. The other mix contained Cas9, two sgRNAs targeting *fhod3b* exons 19 and 20 (of predicted splice isoform X4, Accession XM_068070214, version 1), and ODN for HDR resulting in 2310 bp in-frame deletion in *fhod3b*. Efficacy of mutagenesis was confirmed by genotyping embryos by PCR. Injected embryos were raised to adulthood and crossed to wild-type fish to identify germline founders and generate F1 heterozygotes. Wild-type alleles were identified by PCR. Mutant alleles were subject to Sanger sequence analysis.

The sequences for CRISPR-Cas9 reagents are:

*fhod3a* sgRNA 1 target sequence: GGA CAG CCG AAC ATA GGG GG

*fhod3a* sgRNA 2 target sequence: CGC AGT AAC TCC ATC AAC AT

*fhod3a* ODN: GAC CTT CAG GCT CCG CCC CCT CCT CCC CCA CCC TGC CCC TTT AAC CTC CCG GCT CCG CCC AAC ATC GGC CTG ACA GTT CTT CCT CCG CCG CGC ACC ATC AAA ACT GCC ATC GTC AAC TTC GA

*fhod3b* sgRNA 1 target sequence: TGC GCT GCC CAG CGG AGT GT

*fhod3b* sgRNA 2 target sequence: CAA TAG CTA GCA GAG TCG AA

*fhod3b* ODN: TGA TTC CTA CGG AGG AGG AGA CAC AGA AGA TTC AGG AGG CCC AGC TGG CGA ACC CCG ACA CGA CTC TGC TAG CTA TTG GCA ACT TTC TCA ATG GGA CCA ATG TGA GTG CCA GAA AAG TCT C

Genotyping primers (InVivo Biosystems and Integrated DNA Technologies) used are: forward primer to detect wild-type

*fhod3a*: GCA TGG ACC AAT TAG AAA AG

forward primer to detect *fhod3a* deletion alleles: GAA GGA GTC AGA CTG TAT TTG G

reverse primer for *fhod3a*: TGC TGA GTA GTG TGA GTC TGC

forward primer to detect wild-type *fhod3b*: GCA TGG ACC AAT TAG AAA AG

forward primer to detect *fhod3b* deletion alleles: ATT AGT CAC TGG ATC TGG TCC

reverse primer for *fhod3b*: ATT AGT CAC TGG ATC TGG TCC

All *fhod3a* and *fhod3b* alleles were deletions, although none perfectly matched what was predicted from HDR. *fhod3a^sny3001^* is a deletion of sequence beginning in exon 19 and ending in the following intron (GRCz11, Chr 19: 754,708..756,109). *fhod3a^sny3002^*is an out-of-frame deletion that begins in exon 19 and ends in exon 20 (Chr 19: 754,710..756,307). *fhod3a^sny3003^* is also an out-of-frame deletion that also begins in exon 19 and ends in exon 20 (Chr 19: 754,698..756,304), but with an insertion of “TC” at the deletion site. Assuming the partial intron for *fhod3a^sny3001^* is retained in its mRNA, conceptual translation of all three *fhod3a* alleles produces wild-type Fhod3a through the mid-FH1 domain, followed by 9, 0, or 19 non-Fhod3a amino acids (for *fhod3a^sny3001^*, *fhod3a^sny3002^*, or *fhod3a^sny3003^*, respectively) before a stop codon (Fig.1A). *fhod3b^sny3004^* is a deletion of sequence beginning in the intron between exons 19 and 20, and ending in exon 20 (Chr 16: 18,635,125..18,637,044). Based on a theoretical mRNA for *fhod3b^3004^* that skips exon 20 due to absence of its splice acceptor, a theoretical translation produces wild-type Fhod3b lacking the L helix and M helix of the FH2 domain (Fig.1A). *fhod3b^sny3005^*begins in exon 19 and ends in exon 20, but differs from the in-frame deletion predicted from HDR by absence of 1 bp (Chr 16: 16,635,130..18,637,439). The resulting conceptual translation replaces helix J of the FH2 domain with three non-Fhod3b amino acids followed by a premature stop codon (Fig.1A).

### Fluorescence microscopy

2 dpf and 3 dpf embryos were dechorionated and fixed with 4% PFA overnight at 4°C. 5 dpf animals were anaesthetized with 0.015% tricaine and then fixed in 4% PFA with 0.015% tricaine overnight at 4°C. Embryos were washed with PBST, permeabilized for 8 min with ice-cold acetone, washed in PBST, and blocked for ≥ 1 hr at room temperature with 10% bovine serum albumin (BSA) (Sigma-Aldrich, St. Louis, MO, Cat 3A3059) in PBST. All staining was conducted overnight at 4°C while nutating. Primary antibodies were diluted in 2% BSA/PBST as follows: undiluted for MF 20 (anti-muscle myosin II heavy chain, developed by D. A. Fischman, MD, Cornell University Medical College, obtained from the Developmental Studies Hybridoma Bank (DSHB), created by the NICHD of the NIH and maintained at The University of Iowa, Department of Biology, Iowa City, IA 52242; RRID: AB_2147781); 1:10 for F59 (anti-muscle myosin II heavy chain, developed by F. E. Stockdale, Standford University Medical Center, obtained from the DSHB; RRID: AB_528373); 1:100 for hVIN1 (anti-vinculin, Sigma-Aldrich, Cat #V9264; RRID: AB_477629) and MANDRA1 (anti-dystrophin, Sigma-Aldrich, Cat #D 8043; RRID: AB_2314760); 1:150 for EA5.3 (anti-sarcomeric α-actinin, Invitrogen, Cat #MA1-22863, RRID: AB_557426). Secondary antibody fluorescein conjugated goat anti-mouse (Rockland Immunochemicals, Pottstown, PA, Cat #610-102-121) was diluted 1:200 in 2% BSA/PBST. For immunostained samples, 1:200 diluted Alexa-568-phalloidin (Fisher Scientific, Hanover Park, IL, Cat #A12380) and 4 µg/ml 4’,6-diamidino-2-phenylindole (DAPI) (Fisher Scientific, Hanover Park, IL, Cat #D1306) were included with secondary antibodies. For samples stained with Alexa-568-phalloidin only, phalloidin was diluted into 10%BSA/PBST for incubation.

Stained embryos were mounted in 1.5% low-melt agarose (Sigma-Aldrich, Cat #39346-81-1). Images were acquired on an SP8 laser scanning confocal microscope (Leica, Wetzlar, Germany) driven by LAS X Software (version 3.5.2, build 4758, Leica), and using a 40X/1.25 water immersion objective. Shown fluorescence microscopy images are single confocal slices, except images in Fig.5C showing maximum intensity projections of several confocal slices to simultaneously show fast- and slow-twitch muscle fibers. Fluorescence microscopy images were linearly processed to enhance contrast and were false colored in Adobe Photoshop 2025 (version 26.6.0, Adobe, San Jose, CA).

### Quantification of myofibril damage

To assess myofibril damage in Alexa-568-phalloidin-stained embryos, three confocal sections were imaged for each examined somite. Every muscle fiber visible in each section was manually scored as normal or damaged based on the following criteria. We defined a “myofibril” as any a singular longitudinal F-actin structure with cross striations that spanned a somite from septum to septum. It is likely bundled myofibrils were miscategorized as a single myofibril, particularly in muscle fibers of older embryos where the F-actin content is significantly increased. Thus, our results likely do not reflect the precise proportion of damaged-to- undamaged individual myofibrils within a strain, but this provides a useful metric for comparing different strains of a given age. A myofibril was defined as “damaged” if it was associated with F-actin strands that were oriented obliquely or perpendicularly to the myofibril (as indicated by *arrows,* Fig.4A), or there were obvious discontinuities in the myofibril (as indicated by *arrowheads*, Fig.4A). For 3 dpf and 5 dpf embryos, myofibrils were analyzed in somite 5 and in the penultimate somite of the tail in order to sample muscle fibers of the anterior and the posterior. For 2 dpf embryos, myofibrils were analyzed in somite 5 only, as the muscle fibers in the tail region were too variable in terms of developmental stage. Manual assignment of damaged or undamaged was conducted while blinded to sample identity. Six fish were analyzed for each strain and age, distributed across three different experiments, except for 48 hpf embryos, which were oversampled with 11, 11, 9, and 7 fish for wild-type, *fhod3a^sny3001^*, *fhod3a^sny3002^*, and *fhod3a^sny3003^*, respectively.

### Polarized light microscopy

5 dpf embryos were anesthetized in 0.015% tricaine and pipetted onto glass slides. Excess water was removed for imaging, after which embryos were euthanized. Images were acquired on a Nikon Ti-S microscope (Nikon, Tokyo, Japan), using a Nikon 10x/0.25 Ph1 DL WD 7.0 objective, and a Canon EOS REBEL T3i/EOS 600D digital camera (Canon, Melville, NY) with auto-contrast and auto-focus disabled. For each image, two longitudinal lines were drawn in ImageJ (version 2.0.0-rc-65/1.51g),^57^ one dorsal to and one ventral to the notochord. Each line was drawn to span six somites. Birefringence signal intensity was measured for each line in ImageJ and then plotted in Microsoft Excel for Mac (version 16.91.1; Microsoft Corporation, Redmond, WA).

### Zebrafish injections

To assay the effects of paralysis on *fhod3a* mutant skeletal muscle, mRNA for α-BTX was transcribed from linearized plasmid pmtb-t7-alpha-bungarotoxin (Addgene, Watertown, MA, Cat #69542) with T7 RNA polymerase (Promega, Madison, WI, Cat #P2075), followed by treatment with DNAse I, precipitation with LiCl at −20°C, and ethanol wash. Embryos were injected with 5 nL 25 ng/µL mRNA at the 1- to 2-cell stage, and dechorionated at 24 hpf. Embryos were tested for response to poking at 3 pdf. Those with persistent paralysis were anesthetized with 0.015% tricaine and fixed and processed for F-actin stain.

To assay for disruption to the integrity of the sarcolemma of skeletal muscle fibers, embryos were injected at 5 dpf in the common cardinal vein with 5 nL 0.1% Evans blue dye (Sigma-Aldrich, Cat #E2129, CAS 314-13-6) in Ringer’s solution (155 mM NaCl, 5 mM KCl, 2 mM CaCl_2_, 1 mM MgCl_2_, 2 mM Na_2_HPO_4_, 10 mM HEPES, pH 7.4, 10 mM glucose), as described.^58^ Embryos were allowed to recover 4.75 h, then anesthetized with 0.015% tricaine and live-mounted in 1.5% low-melt agarose for imaging for Evans blue fluorescence on an SP8 laser scanning confocal microscope.

### Statistical analyses

Graphs were made in Prism 10 (version 10.1.1; GraphPad Software, Boston, MA). Data are expressed as average ± SD and individual data points. For experiments where quantitative results were compared, statistical analysis was performed using one-way Analysis of Variation (ANOVA), followed by Tukey’s multiple comparisons *post hoc* test, using Prism 10. Categorical data (Table 1, Table 2) were subject to Chi Squared Tests against the null hypothesis that phenotype occurs independently of homozygosity of the mutant allele, using Microsoft Excel for Mac. *p* < 0.05 was considered statistically significant.

## ACKNOWLEDGMENTS

We thank Hani Suleiman for providing *fhod1^sa12078^* fish. Gene sequence and expression data for this paper were retrieved from the Zebrafish Information Network (ZFIN), University of Oregon, Eugene, OR 97403-5274; URL: http://zfin.org/. This work was supported by a Pilot Grant from the Francis Hendricks Endowment Fund (to D.P. and J.A.) and by R01HD099031 from the Eunice Kennedy Shriver National Institute of Child Health and Human Development (to J.A.). The authors have no conflicts of interest to declare.

## REFERENCES

1. Henderson CA, Gomez CG, Novak SM, Mi-Mi L, Gregorio CC. Overview of the muscle cytoskeleton. Compr Physiol. 2017;7(3):891–944. 10.1002/cphy.c160033

2. Bloch RJ, Gonzalez-Serratos H. Lateral force transmission across costameres in skeletal muscle. Exerc Sport Sci Rev. 2003;31(2):73–78. 10.1097/00003677-200304000-00004

3. Chesarone MA, DuPage AG, Goode BL. Unleashing formins to remodel the actin and microtubule cytoskeletons. Nat Rev Mol Cell Biol. 2010;11(1):62–74. 10.1038/nrm2816

4. Bechtold M, Schultz J, Bogdan S. FHOD proteins in actin dynamics – a formin’ class of its own. Small GTPases. 2014;5(2):11. 10.4161/21541248.2014.973765

5. Kan-O M, Takeya R, Abe T, Kitajima N, Nishida M, Tominaga R, Kurose H, Sumimoto H. Mammalian formin Fhod3 plays an essential role in cardiogenesis by organizing myofibrillogenesis. Biol Open. 2012;1(9):889–896. 10.1242/bio.20121370

6. Taniguchi K, Takeya R, Suetsugu S, Kan-O M, Narusawa M, Shiose A, Tominaga R, Sumimoto H. Mammalian formin fhod3 regulates actin assembly and sarcomere organization in striated muscles. J Biol Chem. 2009;284(43):29873–29881. 10.1074/jbc.m109.059303

7. Iskratsch T, Lange S, Dwyer J, Kho AL, dos Remedios C, Ehler E. Formin follows function: a muscle-specific isoform of FHOD3 is regulated by CK2 phosphorylation and promotes myofibril maintenance. J Cell Biol. 2010;191(6):1159–1172. 10.1083/jcb.201005060

8. Fenix AM, Neininger AC, Taneja N, Hyde K, Visetsouk MR, Garde RJ, Liu B, Nixon BR, Manalo AE, Becker JR, Crawley SW, Bader DM, Tyska MJ, Liu Q, Gutzman JH, Burnette DT. Muscle-specific stress fibers give rise to sarcomeres in cardiomyocytes. Elife. 2018;7:e42144. 10.7554/elife.42144

9. Wooten EC, Hebl VB, Wolf MJ, Greytak SR, Orr NM, Draper I, Calvino JE, Kapur NK, Maron MS, Kullo IJ, Ommen SR, Bos JM, Ackerman MJ, Huggins GS. Formin homology 2 domain containing 3 variants associated with hypertrophic cardiomyopathy. Circ Cardiovasc Genet. 2013;6(1):10–18. 10.1161/circgenetics.112.965277

10. Shwartz A, Dhanyasi N, Schejter ED, Shilo BZ. The *Drosophila* formin Fhos is a primary mediator of sarcomeric thin-filament array assembly. Elife. 2016;5:e16540. 10.7554/elife.16540

11. Kimmich MJ, Sundaramurthy S, Geary MA, Lesanpezeshki L, Yingling CV, Vanapalli SA, Littlefield RS, Pruyne D. FHOD-1 and profilin protect sarcomeres against contraction-induced deformation in *C. elegans*. Mol Biol Cell. 2024;35(11):ar137. 10.1091/mbc.e24-04-0145

12. Pruyne D, Evangelista M, Yang C, Bi E, Zigmond S, Bretscher A, Boone C. Role of formins in actin assembly: nucleation and barbed-end association. Science. 2002;297(5581):612-615. 10.1126/science.1072309

13. Sagot I, Rodal AA, Moseley J, Goode BL, Pellman D. An actin nucleation mechanism mediated by Bni1 and profilin. Nat Cell Biol. 2002;4(8):626–631. 10.1038/ncb834

14. Xu Y, Moseley J, Sagot I, Poy F, Pellman D, Goode BL, Eck MJ. Crystal structures of a Formin Homology-2 domain reveal a tethered dimer architecture. Cell. 2004;116(5):711–723. 10.1016/s0092-8674(04)00210-7

15. Kovar DR, Kuhn JR, Tichy AL, Pollard TD. The fission yeast cytokinesis formin Cdc12p is a barbed end actin filament capping protein gated by profilin. J Cell Biol. 2003;161(5):875–887. 10.1083/jcb.200211078

16. Oosterheert W, Sanders MB, Funk J, Prumbaum D, Raunser S, Bieling P. Molecular mechanism of actin filament elongation by formins. Science. 2024;384(6692):eadn9560. 10.1126/science.adn9560

17. Otomo T, Tomchick DR, Otomo C, Panchal SC, Machius M, Rosen MK. Structural basis of actin filament nucleation and processive capping by a formin homology 2 domain. Nature. 2005;433(7025):488-494. 10.1038/nature03251

18. Thompson ME, Heimsath EG, Gauvin TJ, Higgs HN, Kull FJ. FMNL3 FH2-actin structure gives insight into formin-mediated actin nucleation and elongation. Nat Struct Mol Biol. 2013;20(1):111–118. 10.1038/nsmb.2462

19. Patel AA, Oztug Durer ZA, van Loon AP, Bremer KV, Quinlan ME. *Drosophila* and human FHOD family formin proteins nucleate actin filaments. J Biol Chem. 2018;293(2):532–540. 10.1074/jbc.m117.800888

20. Fujimoto N, Kan-O M, Ushijima T, Kage Y, Tominaga R, Sumimoto H, Takeya R. Transgenic expression of the formin protein Fhod3 selectively in the embryonic heart: role of actin-binding activity of Fhod3 and its sarcomeric localization during myofibrillogenesis. PLoS One. 2016;11(2):e0148472. 10.1371/journal.pone.0148472

21. Pring M, Evangelista M, Boone C, Yang C, Zigmond SH. Mechanism of formin-induced nucleation of actin filaments. Biochemistry. 2003;42(2):486–496. 10.1021/bi026520j

22. Kovar DR, Pollard TD. Insertional assembly of actin filament barbed ends in association with formins produces piconewton forces. Proc Natl Acad Sci U S A. 2004;101(41):147250–14730. 10.1073/pnas.0405902101

23. Valencia DA, Koeberlein AN, Nakano H, Rudas A, Harui A, Spencer C, Nakano A, Quinlan ME. Human formin FHOD3-mediated actin elongation is required for sarcomere integrity in cardiomyocytes. bioRxiv [Preprint]. 2024;10.13.618125. 10.1101/2024.10.13.618125

24. Gore AV, Monzo K, Cha YR, Pan W, Weinstein BM. Vascular development in the zebrafish. Cold Spring Harb Perspect Med. 2012;2(5):a006684. 10.1101/cshperspect.a006684

25. Stickney HL, Barresi MJ, Devoto SH. Somite development in zebrafish. Dev Dyn. 2000;219(3):287–303. 10.1002/1097-0177(2000)9999:9999%3C::aid-dvdy1065%3E3.0.co;2-a

26. Thisse B, Thisse C. Fast release clones: a high throughput expression analysis. ZFIN Direct Data Submission. 2004;http://zfin.org.

27. Farnsworth DR, Saunders LM, Miller AC. A single-cell transcriptome atlas for zebrafish development. Dev Biol. 2020;459(2):100–108. 10.1016/j.ydbio.2019.11.008

28. Sur A, Wang Y, Capar P, Margolin G, Prochaska MK, Farrell JA. Single-cell analysis of shared signatures and transcriptional diversity during zebrafish development. Dev Cell. 2023;58(24):3028–3047.e12. 10.1016/j.devcel.2023.11.001

29. Devoto SH, Melançon E, Eisen JS, Westerfield M. Identification of separate slow and fast muscle precursor cells *in vivo*, prior to somite formation. Development. 1996;122(11):3371–3380. 10.1242/dev.122.11.3371

30. Schapira G, Dreyfus JC, Joly M. Changes in the flow birefringence of myosin as a result of muscular atrophy. Nature. 1952;170(4325):494-495. 10.1038/170494b0

31. Mercuri E, Bönnemann CG, Muntoni F. Muscular dystrophies. Lancet. 2019;394(10213):2025-2038. 10.1016/s0140-6736(19)32910-1

32. Gibbs EM, Horstick EJ, Dowling JJ. Swimming into prominence: the zebrafish as a valuable tool for studying human myopathies and muscular dystrophies. FEBS J. 2013;280(17):4187–4197. 10.1111/febs.12412

33. Tesoriero C, Greco F, Cannone E, Ghirotto F, Facchinello N, Schiavone M, Vettori A. Modeling human muscular dystrophies in zebrafish: mutant lines, transgenic fluorescent biosensors, and phenotyping assays. Int J Mol Sci. 2023;24(9):8314. 10.3390/ijms24098314

34. Etard C, Behra M, Ertzer R, Fischer N, Jesuthasan S, Blader P, Geisler R, Strähle U. Mutation in the δ-subunit of the nAChR suppresses the muscle defects caused by lack of Dytrophin. Dev Dyn. 2005;234(4):1016–1025. 10.1002/dvdy.20592

35. Berger J, Berger S, Hall TE, Lieschke GJ, Currie PD. Dystrophin-deficient zebrafish feature aspects of the Duchenne muscular dystrophy pathology. Neuromuscul Disord. 2010;20(12):826–832. 10.1016/j.nmd.2010.08.004

36. Li M, Arner A. Immobilization of Dystrophin and Laminin α2-chain deficient zebrafish larvae *in vivo* prevents the development of muscular dystrophy. PLoS One. 2015;10(11):e0139483. 10.1371/journal.pone.0139483

37. Swinburne IA, Mosaliganti KR, Green AA, Megason SG. Improved long-term imaging of embryos with genetically encoded α-bungarotoxin. PLoS One. 2015;10(8):e0134005. 10.1371/journal.pone.0134005

38. Matsuda R, Nishikawa A, Tanaka H. Visualization of dystrophic muscle fibers in *mdx* mouse by vital staining with Evans blue: evidence of apoptosis in dystrophin-deficient muscle. J Biochem. 1995;118(5):959–964. 10.1093/jb/118.5.959

39. Straub V, Rafael JA, Chamberlain JS, Campbell KP. Animal models for muscular dystrophy show different patterns of sarcolemmal disruption. J Cell Biol. 1997;139(2):375–385. 10.1083/jcb.139.2.375

40. Bassett DI, Bryson-Richardson RJ, Daggett DF, Gautier P, Keenan DG, Currie PD. Dystrophin is required for the formation of stable muscle attachments in the zebrafish embryo. Development. 2003;130(23):5851–5860. 10.1242/dev.00799

41. Jaka O, Casas-Fraile L, de Munain AL, Sáenz A. Costamere proteins and their involvement in myopathic processes. Expert Rev Mol Med. 2015;17:e12. 10.1017/erm.2015.9

42. Gut P, Reischauer S, Stainier DYR, Arnaout R. Little fish, big data: zebrafish as a model for cardiovascular and metabolic disease. Physiol Rev. 2017;97(3):889–938. 10.1152/physrev.00038.2016

43. Krainer EC, Ouderkirk JL, Miller EW, Miller MR, Mersich AT, Blystone SD. The multiplicity of human formins: expression patterns in cells and tissues. Cytoskeleton (Hoboken). 2013;70(8):424–438. 10.1002/cm.21113

44. Sanematsu F, Kanai A, Ushijima T, Shiraishi A, Abe T, Kage Y, Sumimoto H, Takeya R. Fhod1, an actin-organizing formin protein, is dispensable for cardiac development and function in mice. Cytoskeleton (Hoboken). 2019;76(2):219–229. 10.1002/cm.21523

45. Sulistomo HW, Nemoto T, Yanagita T, Takeya R. Formin homology 2 domain-containing 3 (Fhod3) controls neural plate morphogenesis in mouse cranial neurulation by regulating multidirectional apical constriction. J Biol Chem. 2019;294(*8*):2924–2934. 10.1074/jbc.ra118.005471

46. Lesurf R, Said A, Akinrinade O, Breckpot J, Delfosse K, Liu T, Yao R, Persad G, McKenna F, Noche RR, Oliveros W, Mattioli K, Shah S, Miron A, Yang Q, Meng G, et al. Whole genome sequencing delineates regulatory, copy number, and cryptic splice variants in early onset cardiomyopathy. NPJ Genom Med. 2022;7(1):18. 10.1038/s41525-022-00288-y

47. Sanger JW, Wang J, Holloway B, Du A, Sanger JM. Myofibrillogenesis in skeletal muscle cells in zebrafish. Cell Motil Cytoskeleton. 2009;66(8):556–566. 10.1002/cm.20365

48. Straub V, Campbell KP. Muscular dystrophies and the dystrophin-glycoprotein complex. Curr Opin Neurol. 1997;10(2):168–175. 10.1097/00019052-199704000-00016

49. Nakagawa H, Kage Y, Miura A, Sulistomo HW, Matsuyama S, Yamashita Y, Takeya R. The expression of the formin Fhod3 in mouse tongue straited muscle. Cell Struct Funct. 2024;49(2):111–122. 10.1247/csf.24044

50. Ushijima T, Fujimoto N, Matsuyama S, Kan-O M, Kiyonari H, Shioi G, Kage Y, Yamasaki S, Takeya R, Sumimoto H. The actin-organizing formin protein Fhod3 is required for postnatal development and functional maintenance of the adult heart in mice. J Biol Chem. 2018;293(1):148–162. 10.1074/jbc.m117.813931

51. Petrany MJ, Swoboda CO, Sun C, Chetal K, Chen X, Weirauch MT, Salomonis N, Millay DP. Single-nucleus RNA-seq identifies transcriptional heterogeneity in multinucleated skeletal myofibers. Nat Commun. 2020;11(1):6374. 10.1038/s41467-020-20063-w

52. Craig SW, Pardo JV. Gamma actin, spectrin, and intermediate filament proteins colocalize with vinculin at costameres, myofibril-to-sarcolemma attachment sites. Cell Motil. 1983;3(5- 6):449–462. 10.1002/cm.970030513

53. Gokhin DS, Fowler VM. Cytoplasmic γ-actin and tropomodulin isoforms link to the sarcoplasmic reticulum in skeletal muscle fibers. J Cell Biol. 2011;194(1):105–120. 10.1083/jcb.201011128

54. Sonnemann KJ, Fitzsimons DP, Patel JF, Liu Y, Schneider MF, Moss RL, Ervasti JM. Cytoplasmic γ-actin is not required for skeletal muscle development but its absence leads to a progressive myopathy. Dev Cell. 2006;11(3):387–397. 10.1016/j.devcel.2006.07.001

55. Prins KW, Lowe DA, Ervasti JM. Skeletal muscle-specific ablation of γ_cyto_-actin does not exacerbate the *mdx* phenotype. PLoS One. 2008;3(6):e2419. 10.1371/journal.pone.0002419

56. Roman W, Martins JP, Carvalho FA, Voituriez R, Abella JVG, Santos NC, Cadot B, Way M, Gomes ER. Myofibril contraction and crosslinking drive nuclear movement to the periphery in skeletal muscle. Nat Cell Biol. 2017;19(10):1189–1201. 10.1038/ncb3605

57. Schneider CA, Rasband WS, Eliceiri KW. NIH image to ImageJ: 25 years of image analysis. Nat Methods. 2012;9(7):671–675. 10.1038/nmeth.2089

58. Smith SJ, Horstick EJ, Davidson AE, Dowling J. Analysis of zebrafish larvae skeletal muscle integrity with Evans blue dye. J Vis Exp. 2015;(105):53183. 10.3791/53183

